# Contrastive pre-training for sequence based genomics models

**DOI:** 10.1101/2024.06.10.598319

**Authors:** Ksenia Sokolova, Kathleen M. Chen, Olga G. Troyanskaya

**Author notes:** Equal contribution (Ksenia Sokolova, Kathleen M. Chen).

## Abstract

**Motivation:** In recent years deep learning has become one of the central approaches in a number of applications, including many tasks in genomics. However, as models grow in depth and complexity, they either require more data or a strategic initialization technique to improve performance.

**Results:** In this project, we introduce cGen, a novel unsupervised, model-agnostic contrastive pretraining method for sequence-based models. cGen can be used before training to initialize weights, reducing the size of the dataset needed. It works through learning the intrinsic features of the reference genome and makes no assumptions on the underlying structure. We show that the embeddings produced by the unsupervised model are already informative for gene expression prediction and that the sequence features provide a meaningful clustering. We demonstrate that cGen improves model performance in various sequence-based deep learning applications, such as chromatin profiling prediction and gene expression. Our findings suggest that using cGen, particularly in areas constrained by data availability, could improve the performance of deep learning genomic models without the need to modify the model architecture.

**Availability and implementation:** Source code available at github.com/ksenia007/cGen

## 1 Introduction

In recent years deep learning has become one of the central approaches in a number of applications, from those in computer vision (CV), to natural language processing (NLP), to genomics [1–4]. However, as models continue to grow in depth and complexity, they either require more data or a strategic initialization technique (or both) to improve performance. Pre-training is one key technique which involves training the model first on some simple but well defined task that has abundant training data, followed by fine-tuning: applying the pre-trained model to the task of interest that has much less training data available [5–7]. The best pre-training tasks are usually ones that are free or cheap to create the datasets for, with unsupervised models taking the lead in NLP and CV [7,8]. However, designing the pre-training task such that it is both unsupervised and useful for the downstream application is not trivial.

In genomics, unsupervised pre-training is an emerging area of research. Borrowing from NLP, masked token prediction and next-token prediction have been used to pre-train models for genomic tasks [9–11]. However, masked-token prediction makes assumptions on the underlying structure of the data, where the genome must be split into “words” - sequences of 1-6bp. While it is possible for the coding genome, where every three nucleotides will form a triplet encoding an amino acid, in non-coding regions of the genome these splits are usually performed randomly. However, the vast majority of pre-training data comes from noncoding sequences that constitute 98% of the human genome. On the other hand, the task of the next token (base pair) prediction, while common in NLP, does not have a straightforward extension to genomics, where there would only 4 possible tokens for single base pair “words” (compared to GPT-2 with vocabulary greater than 50K [12]). Therefore, there is an interest in exploring different pre-training approaches better suited for genomic sequence modeling.

Contrastive pre-training is a specific pre-training approach that has been shown to be highly effective in computer vision. In particular, contrastive pre-training has been shown to improve performance on tasks that benefit from defining the relative embeddings of training points. At the core of the method lies the understanding that similar inputs should be placed near each other in the embedding space, while dissimilar ones are placed further apart. While popular in computer vision, there has been a very limited use of contrastive models in genomics, focused on relatively simple tasks such as motif occupancy [13].

In this paper we introduce cGen, a novel model-agnostic contrastive pre-training framework for genomic sequences modeling. cGen is inspired by the simCLR [14][15] method popular in computer vision, where pairs of augmented versions of the same image are used as positive samples and other images as negatives. cGen creates negative and positive pairs through augmentation of randomly sampled genomic sequences. Through systematic analysis of genomics-specific augmentations, we discover the best training augmentations, and show improvements in performance on downstream genomics tasks. Overall, we find that cGen unsupervised pre-training produces meaningful embeddings that are informative of regulatory features of the sequence and gene expression.

## 2 Methods

cGen architecture: Our model had two modules: a “core” encoder block, that accepts the one hot encoded sequence, and a fully connected head. The encoder block is model agnostic, and in this project we used architectures similar to [2,16]. These represented different sizes of the CNN, where Sei [2] was a large CNN based model with skip connections. We modify [16] to accommodate for the expanded sequence input by adding more convolutional blocks into the architecture and changing the fully connected layer (Supp. Table S1). Sei is modified only in the fully connected head to fit into our framework.

Training details: PytorchLightning[17] framework with Pytorch was used for the project. We use 500 batches in training, and run for a maximum of 100 epochs. Model with the best validation loss is used unless otherwise noted. We performed a weak search of the learning rate, where we evaluated starting learning rates [0.1, 0.01, 0.001, 0.0001, 0.00001]. For optimizers, we used AdamW for [16] and SGD with momentum 0.9 for [2]. The fully connected head is changed depending on the task in the finetuning. In pre-training, this layer is used for projection, and is discarded when training the final model. We use the mlp size of 256 or 512 for the contrastive pre-training independent of the model.

Linear probes: For linear probing, we used GTEx tissue-level gene expression data with architecture from [18]. To reduce overfitting and increase the training size of the probe, when compiling the dataset, we created multiple entries for each gene, where the expression stays the same but the input sequence is shifted around TSS +/-10bp in 1 bp increments. That is done to expand the dataset, and prevent model memorization, forcing the model to learn features of the sequence. We log-transformed gene expression values w/ the pseudocount of 0.1, similar to [16,19]. LinearRegression implementation from scikit package was used for each of the tissues separately, and spearman score was used from the scipy.stats package.

Pre-training approaches: For the pre-train, we passed in augmented sequences and used InfoNCE loss [20]. When fine tuning, we replaced the fully connected head with new randomly initialized layers (as recommended in [14]). The search for augmentation and other large-scale benchmarking was performed on the [16], and then reproduced on the larger Sei architecture. The primary genomics-specific augmentations in cGen include:

- Flip sequence: Flipping the sequence left to right + make a complement.
- Masking up to a % of the sequence base pairs (0%, 10%, 20%, 30%, 40%). For every sequence the exact number masked is random.
- Shifts left or right by maximum of N base pairs (Shift of 1, 100, 400). For each of the samples in augmentations, numpy randint() is used to draw a specific number used.

Gene expression finetuning: To evaluate augmentations, we used 10% of the training samples and compared the MSE loss (torch.nn) on the validation set. Train/validation split was the same between the models. In addition, we used a starting learning rate of 0.01, weight decay 0.0000001, batch size 64, sequence length 4096 (Supp. Tables S1-2).

Genomic benchmarks finetuning: The genomic benchmarks datasets were downloaded using the genomic-benchmarks (version 0.0.9) [21] Python package. We used their pre-specified training, validation, and test sets. Models were trained on the enhancer prediction tasks at learning rate of 0.001, weight decay 0.0000001, batch size 64. Sequence length was pre-specified for each individual task.

Chromatin profile finetuning: The 2,002 chromatin profiles encompass transcription factors, histone marks, and chromatin accessibility from ENCODE [22] and Roadmap Epigenomics [23] projects, and the dataset matches that in [16]. We use chromosomal holdouts for validation (chr10) and testing (chr8, chr9), learning rate of 0.01, weight decay 0.0000001, batch size 64, sequence length 4096.

## 3 Results

We developed a flexible model-agnostic contrastive pre-training approach for genomics sequence models. It is a pre-training step that can be used with CNN, transformer or other architectures. At its core, the model is pre-trained by learning to differentiate the same sequence that has been augmented, against other independently sampled sequences from the genome (Figure 1). The goal is for the model to fill in the gaps of the augmentations without forcing it to memorize specific base pairs.

**Figure 1:**
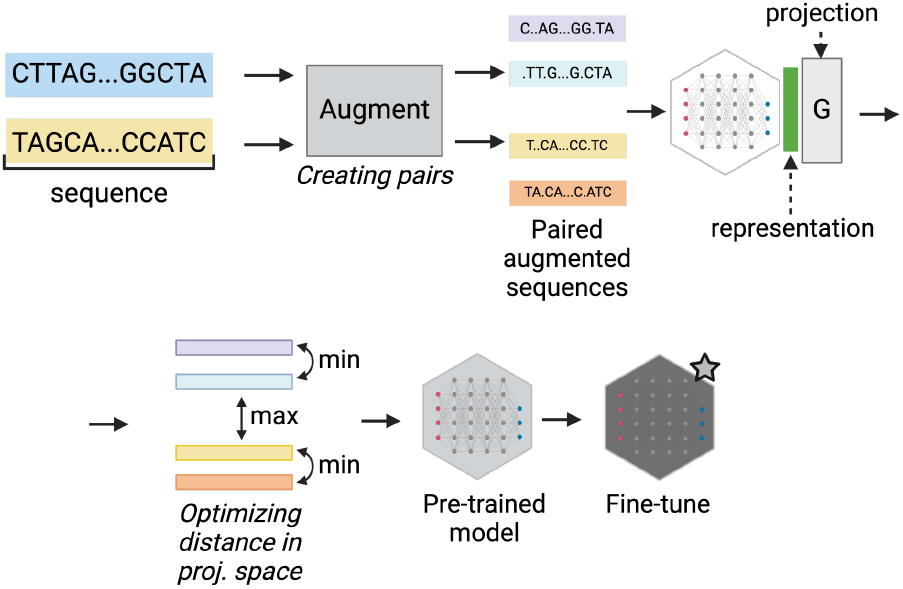
Contrastive pre-training overview. In the pre-training step, the inputs in the model are sequences drawn i.i.d. from the reference genome. Each sequence is augmented to create an alternative (paired) view and passed through the network. The goal at the pre-training step is optimizing the distances between resulting embeddings – that is, the distance between paired sequences is minimized in the embedding space, while distances to other sequence samples are maximized. The pre-trained model is then used in the fine-tuning step where limited data is available, resulting in a context-specific model.

Specifically, the sequences used for pre-training are sampled iid from the reference genome, where each sequence is augmented to create a matched pair. It is then passed through the model, followed by a projection head [14], which maps the internal representation into the space used for contrastive loss.

To approximate the information content achieved by using contrastive pre-training alone, we used linear classifier probes [24] on top of the representation embedding with input length of 10500. This approach allows us to measure how much biologically relevant information is learnt during pre-training. The linear probing was applied to the prediction of tissue-level gene expression collected by GTEx, a complex task that requires sequence context understanding (Figure 2).

**Figure 2:**
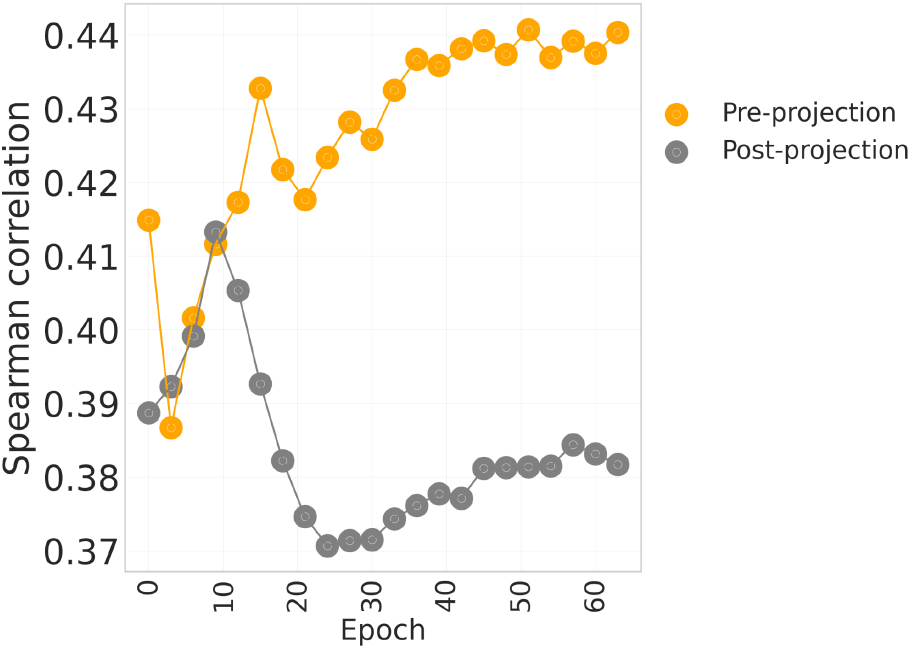
Mean of the performances across GTEx tissues over pre-training epochs. Performance across the tissues deteriorates if the embedding is taken after the projection.

We observe that the embeddings are informative. A random input model predicts a constant value, while a pre-trained model achieves a mean Spearman’s rank correlation of 0.44 across the tissues. There is a clear trend toward improving scores as the model trains (Figure 2). Scores after the first epoch are significantly lower than the scores after the 63rd epoch (p-value=3.74e-08), as we would expect from informative embeddings. In addition, only the “pre-projection” scores are informative, and using the score after the projection in the loss space diminishes the linear probe performance (Figure 2). This is likely due to the loss of information as a result of projection head optimizing for the InfoNCE loss (for example it should be invariant to transformation), and is in agreement with previous work on the contrastive pre-training in computer vision [14].

Further, the unsupervised embeddings exhibit clustering patterns that align with known genome features. We visualized the embeddings on randomly sampled ∼200k sequences from the human reference genome using UMAP [25] and colored each point based on its corresponding regulatory sequence class assignment [2] (Figure 3A). The most pronounced clusters in this visualization show a strong division between regulatory sequences and repeat regions (Centromere, Heterochromatin). Promoter sequences in particular are strongly clustered.

**Figure 3:**
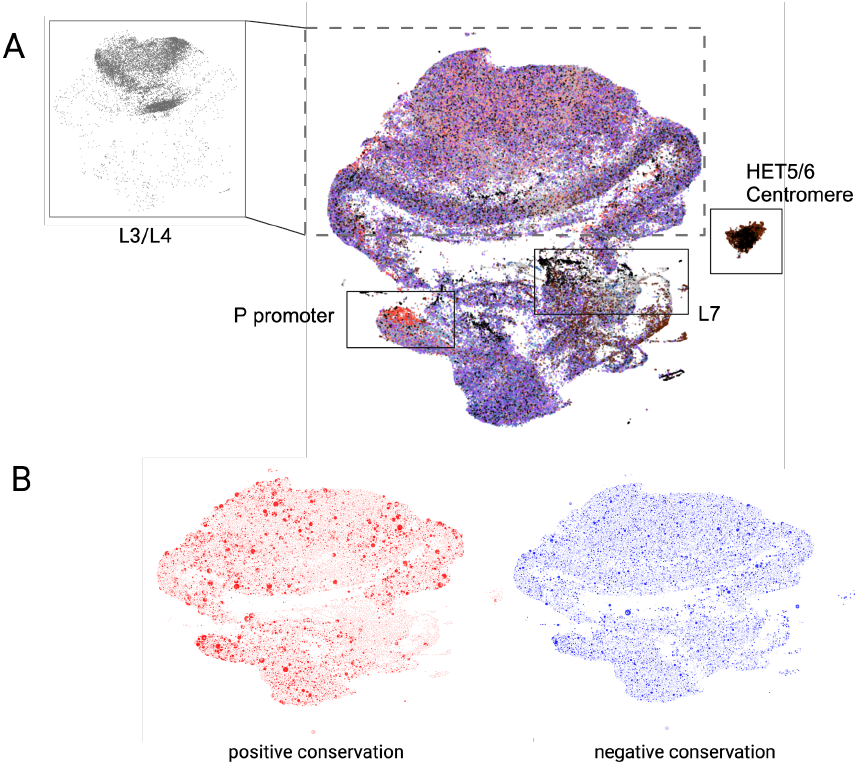
Clustering of the unsupervised embeddings. (A) Showing embeddings colored by the associated sequence class, as defined by [2]. Without any labeled information, meaningful clusters start forming. (B) Embeddings can be separated based on the sign of the PhyloP [26] evolutionary constraint of the sequence, the size of the dot corresponds to the magnitude of the score. No strong clustering based on the conservation scores is observed, apart from the expected enrichment of the positive conserved scores in the area of the P-promoter sequences.

This clustering was obtained in an unsupervised manner, meaning that the model has learnt clusters from the sequence augmentation strategy specified during pre-train alone. Importantly, we do not observe clustering connected to the conservation scores, apart from the expected enrichment of the positive conserved scores in the regulatory promoter sequences (Figure 4B).

**Figure 4:**
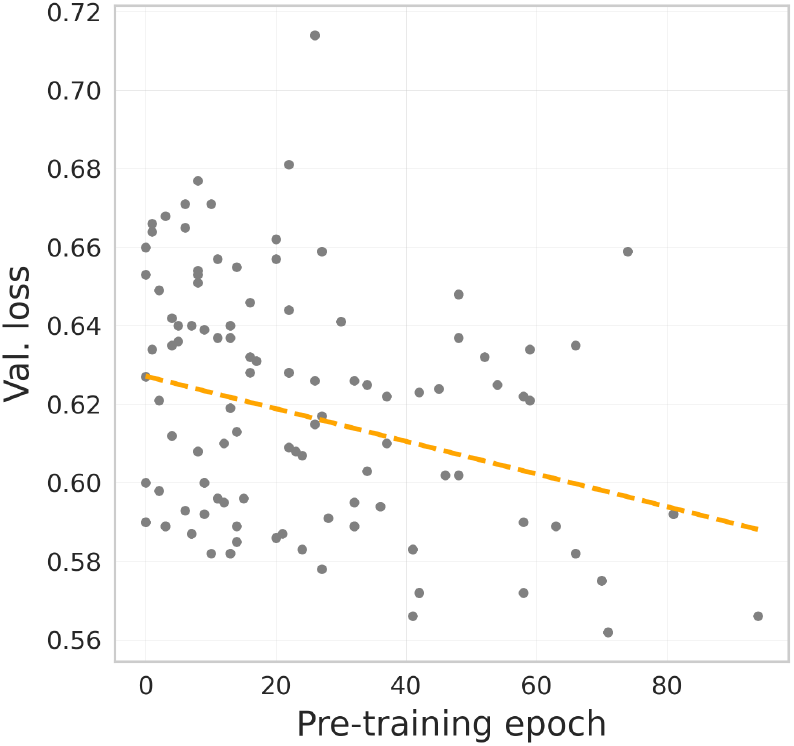
Training for more epochs improves model performance. Validation loss of the fine-tuned model was decreasing when the pre-training model was trained longer. Showing results for the pre-trained model with 40% masking augmentation.

Contrastive pre-training relies heavily on the construction of relevant augmentations to create pairs. While in the previous section we used a manually selected set of augmentations, our goal was to discover tasks that would result in highest gains for downstream prediction of complicated context-specific features in a consistent manner.

To do so, we define a set of sequence-based augmentations, and check pairwise combinations of these augmentations for changes to downstream performance. Specifically, we used a tissue-independent gene expression dataset from [18], and in the fine-tuning use only 10% of the dataset. We observe a strong trend toward more complicated augmentation tasks improving the performance (Table 1). Easy tasks such as flipping the sequence with 50% probability or shifting just one base pair did not lead to any improvement and could worsen the downstream performance. Masking 20% of the base pairs or higher, coupled with random flips of the sequence, leads to the highest gains in performance, closely followed by strong shifts.

**Table 1:** Validation loss is used as a measure of the model’s performance, with % improvement over the non-pretrained model. All of the models were trained on the 10% of available data and with 256 batch size for a maximum of 100 epochs. Results for the best epoch are shown and colored based on extent of improvement (yellow to green). See Supp. Table 2 for values.

|  | Flip 50%<br>chance | Shift 100 | Shift 400 | Shift 1 |
| --- | --- | --- | --- | --- |
| Flip 50% | -3.31% |  |  |  |
| Shift 100 | 6.78% | 10.73% |  |  |
| Shift 400 | 2.68% |  | -3.79% |  |
| Shift 1 | -3.31% |  |  | 1.89% |
| Mask 10% | -0.32% | 8.99% | 1.74% | 0.32% |
| Mask 20% | 12.30% | 3.79% | 6.94% | 2.05% |
| Mask 30% | 9.94% | 9.46% | 7.73% | 7.41% |
| Mask 40% | 9.46% | 10.73% | 2.05% | 8.52% |

Another parameter critical for model pre-training is the number of epochs needed. We thus investigated whether longer pre-training leads to model improvement or overfits to the task. We selected a “hard” pretrain of 40% masking and performed fine tuning for not only for the best pre-trained model, but also for the earlier epochs. In agreement with the linear probing, longer pre-training results in better results, with some fluctuation in the early epochs (Figure 4).

We observed that training using cGen improved performance of the models. For the gene expression task, the improvement was sustained across various amounts of training data used (Table 2). Note that these results are for predicting gene expression from a short sequence (4096bp), with a CNN architecture similar to [16] (updated for 4096 sequence length), and are not directly comparable to the state-of-the-art gene expression models. However, they clearly demonstrate the importance of pre-training to downstream performance. The results are sustained for the larger Sei model, where, on the 10% of the gene expression and same sequence length, the model achieves similar results (r2=0.381 for training from scratch and r2=0.420 with pre-training).

**Table 2:** Effect of pre-training on performance. Pre-trained model outperformed no pretrain on all data subsets. Architecture used is adapted from [16], see Supp.Table 1.

| Fraction data | R2 values |  | Loss |  |
| --- | --- | --- | --- | --- |
|  | no pretrain | w/ pretrain | no pretrain | w/ pretrain |
| 1% | 0.33478 | **0.3706** | 0.677 | **0.643** |
| 5% | 0.3524 | **0.3881** | 0.644 | **0.626** |
| 10% | 0.3599 | **0.42963** | 0.634 | **0.566** |
| 100% | 0.4191 | **0.5458** | 0.566 | **0.442** |

To more comprehensively evaluate the efficacy of the cGen pre-training framework, we fine-tuned a pre-trained Sei model with 40% masking augmentation on additional tasks. For the chromatin profiling task of 2,002 chromatin profiles [16,27], we show that fine-tuned Sei achieves an average AUC of 0.855, while Sei trained from scratch has an average AUC of 0.853 on the test holdout dataset (50 epochs of training). We further fine tune on the two enhancer prediction tasks from Genomic Benchmarks (human enhancers Cohn et al., human enhancers Ensembl) [21]. Fine-tuned Sei achieves a top-1 accuracy of 73.1% on the human enhancers Cohn task, whereas Sei trained from scratch achieves a top-1 accuracy of 72.3%. For human enhancers Ensembl, fine-tuned Sei achieves top-1 accuracy of 88.4%, but Sei trained from scratch on the same parameters was not able to converge with the given training parameters. This illustrates another possible benefit of pre-training: the pre-trained weight initialization can accelerate training over multiple tasks. The performance reported by Sei does in fact surpass the benchmark performance by CNN stated in [21] and is on-par with current SOTA model performance [28].

## 5 Conclusions and discussion

We developed a genomics-specific contrastive pre-training framework, cGen, that uses genomics-specific augmentations and pretraining to improve performance on downstream tasks. By focusing on relationships between different inputs, contrastive pre-training allowed the model to learn a meaningful representation of the underlying genomic data, which can then be fine-tuned for specific downstream tasks. The self-supervised embeddings were informative of the underlying genomic classes without any supervised input. Importantly, this pre-training is not task specific, and as such it could be used for any sequence-based problem, ranging from variant imputation to sequence classification.

Harder pre-training tasks resulted in better fine tuning performance. In the future, it could be helpful to incorporate more biological knowledge into the generation of alternative views in the augmentation, for example guided by evolutionary constraints. In addition, we have found that some pre-trains did not improve performance, most likely due to the randomness of training. While pre-training may not invariably lead to substantial improvements, it could enhance the generalization capabilities of models and their resilience to perturbations [29]. In the future, it could be of interest to explore whether this holds true for cGen.

## Acknowledgements

The authors acknowledge all members of the Troyanskaya laboratory at Princeton University and the Flatiron Institute for helpful discussions. We also thank the Simons Foundation and the Scientific Computing Core of the Flatiron Institute. We are pleased to acknowledge that the work reported on in this paper was substantially performed using the Princeton Research Computing resources at Princeton University which is a consortium of groups led by the Princeton Institute for Computational Science and Engineering (PICSciE) and Office of Information Technology’s Research Computing.

