## Supplemental file for "Contrastive pre-training for sequence based genomics models"

### Supplementary materials

```
(backbone): ModelContrastive(
  (backbone): ModelBody(
    (backbone): Sequential(
      (0): Conv2d(4, 320, kernel_size=(1, 8), stride=(1, 1))
      (1): ReLU()
      (2): Conv2d(320, 320, kernel_size=(1, 8), stride=(1, 1))
      (3): ReLU()
      (4): Dropout(p=0.2, inplace=False)
      (5): MaxPool2d(kernel_size=(1, 4), stride=(1, 4), padding=0, dilation=1, ceil_mode=False)
      (6): Conv2d(320, 480, kernel_size=(1, 8), stride=(1, 1))
      (7): ReLU()
      (8): Conv2d(480, 480, kernel_size=(1, 8), stride=(1, 1))
      (9): ReLU()
      (10): Dropout(p=0.2, inplace=False)
      (11): MaxPool2d(kernel_size=(1, 4), stride=(1, 4), padding=0, dilation=1, ceil_mode=False)
      (12): Conv2d(480, 640, kernel_size=(1, 8), stride=(1, 1))
      (13): ReLU()
      (14): Conv2d(640, 640, kernel_size=(1, 8), stride=(1, 1))
      (15): ReLU()
      (16): MaxPool2d(kernel_size=(1, 4), stride=(1, 4), padding=0, dilation=1, ceil_mode=False)
      (17): Conv2d(640, 640, kernel_size=(1, 8), stride=(1, 1))
      (18): ReLU()
      (19): Conv2d(640, 640, kernel_size=(1, 8), stride=(1, 1))
      (20): ReLU()
      (21): MaxPool2d(kernel_size=(1, 4), stride=(1, 4), padding=0, dilation=1, ceil_mode=False)
      (22): Conv2d(640, 640, kernel_size=(1, 4), stride=(1, 1))
      (23): ReLU()
    )
  )
  (contr_layer): Sequential(
    (0): Linear(in_features=5120, out_features=256, bias=True)
    (1): ReLU(inplace=True)
    (2): Linear(in_features=256, out_features=256, bias=True)
  )
)
```

*Supplementary Table 1: Architecture used in pre-training, related to table 1. The model accepts a sequence input of length 4096. In fine-tuning the Sequential() block is replaced.*

| *Pre-training augmentations* | | | *Finetuning* |
|---|---|---|---|
| **Shift** | **Mask** | **Flip / prob** | **Loss** |
| 1 | 0.2 | T / 0.5 | 0.556 |
| 100 | 0 | F | 0.566 |
| 100 | 0.4 | F | 0.566 |
| 1 | 0.3 | T / 0.5 | 0.571 |
| 1 | 0.4 | T / 0.5 | 0.574 |
| 100 | 0.3 | F | 0.574 |
| 100 | 0.1 | F | 0.577 |
| 1 | 0.4 | F | 0.58 |
| 400 | 0.3 | F | 0.585 |
| 1 | 0.3 | F | 0.587 |
| 400 | 0.2 | F | 0.59 |
| 100 | 0 | T / 0.5 | 0.591 |
| 100 | 0 | T / 0.5 | 0.594 |
| 100 | 0.2 | F | 0.61 |
| 400 | 0 | T / 0.5 | 0.617 |
| 1 | 0.2 | F | 0.621 |
| 400 | 0.4 | F | 0.621 |
| 1 | 0 | F | 0.622 |
| 400 | 0.1 | F | 0.623 |
| 1 | 0.1 | F | 0.632 |
| 1 | 0.1 | T / 0.5 | 0.636 |
| 1 | 0 | T / 0.5 | 0.655 |
| 400 | 0 | F | 0.658 |

*Supplementary Table 2: Augmentations combinations tried and resulting scores. Showing results related to Table 1. All the models had the same batch size (256), and same training parameters in both training and finetuning.*
